# Hypothalamic dopamine neurons control sensorimotor behavior by modulating brainstem premotor nuclei

**DOI:** 10.1101/2020.07.13.195982

**Authors:** Joshua P. Barrios, Wei-Chun Wang, Roman England, Erica Reifenberg, Adam D. Douglass

## Abstract

Dopamine (DA)-producing neurons are critically involved in the production of motor behaviors in multiple circuits that are conserved from basal vertebrates to mammals. While there is increasing evidence that DA neurons in the hypothalamus play a locomotor role, their precise contributions to behavior and the circuit mechanisms by which they are achieved remain unclear. Here we demonstrate that *tyrosine hydroxylase 2*-expressing (*th2*+) DA neurons in the zebrafish hypothalamus fire phasic bursts of activity to acutely promote swimming and modulate audiomotor behaviors on fast timescales. Their anatomy and physiology reveal two distinct functional DA modules within the hypothalamus. The first comprises an interconnected set of cerebrospinal fluid-contacting DA nuclei surrounding the third ventricle, which lack distal projections outside of the hypothalamus and influence locomotion through unknown means. The second includes neurons in the preoptic nucleus, which send long-range projections to targets throughout the brain, including the mid- and hindbrain, where they activate premotor circuits involved in swimming and sensorimotor integration. These data suggest a broad regulation of motor behavior by DA neurons within multiple hypothalamic nuclei and elucidate a novel functional mechanism for the preoptic DA neurons in the initiation of movement.

## Introduction

Vertebrates are capable of an especially complex locomotor repertoire, with a variety of adaptive behaviors intended to address specific environmental or internal demands and each requiring the recruitment of specific neural circuits. The modulatory neurotransmitter dopamine (DA) plays an essential role in this process, being involved in the production, selection, and modulation of motor behaviors in multiple circuits contexts including the basal ganglia[1, 2] and spinal cord[3–5]. Dopamine-producing neurons also exist in five regions of the hypothalamus[6–8] (A11-A15 groups), and a number of anatomical, pharmacological, and lesion studies have implicated these areas in the production of a variety of locomotor behaviors[6, 9–11], yet their precise behavioral functions and the circuit mechanisms involved are not well understood.

The larval zebrafish has become a key tool in the study of the structure and function of neural circuits due to its genetic tractability, optical transparency, and simplified vertebrate anatomy[12, 13]. While most structures in the zebrafish brain appear to be highly conserved in mammals[14, 15], a thorough identification of the anatomical and functional homologs of specific DA nuclei has been hampered by a lack of connectivity information and functional data. This is particularly true for neurons expressing *tyrosine hydroxylase 2* (*th2*)[14]. This second isoform of *tyrosine hydroxylase*, the rate-limiting enzyme in DA production, was discovered relatively recently and is present in all non-mammalian vertebrates[16, 17]. The development of antibodies[18] and genetic lines[19] has very recently made it possible to begin to study the anatomy and function of these neurons. Unlike *th1*, which is broadly expressed in several brain regions[20], *th2* is expressed only in four hypothalamic nuclei: the preoptic nucleus (PON), the posterior tuberculum (PT), and the intermediate (Hi), and caudal hypothalamus (Hc)[18, 20]. The latter three of these groups are distinct in being closely associated with the 3^rd^ ventricle and in containing neurons that directly contact the cerebrospinal fluid (“liquor-contacting neurons”, LCNs;[20, 21]). All four groups show little co-expression of *th1*, making the *th2* enhancer a useful and specific tool for studying hypothalamic sources of DA.

Previous work from our lab has shown that the *th2*+ neurons are necessary to maintain normal levels of spontaneous locomotor activity[19], and that activating the entire genetically-defined cohort of these cells elicits an increase in movement. Which of the four *th2+* nuclei mediate these effects, and the functional targets through which they operate, are unknown. The temporal relationship between their activity and behavior is similarly unclear. Recent work has shown that phasic release of DA in the striatum can acutely modulate behavior on much faster timescales than previously thought and that distinct DAergic axons within a single anatomical projection either modulate motor behavior or signal reward[22, 23]. It remains to be seen if this fast action and functional heterogeneity are common features of other DAergic clusters, such as those in the hypothalamus.

Here, we show that hypothalamic *th2*+ neurons promote a variety of locomotor behaviors in a phasic and intensity-dependent manner. A subset of these cells also responds to acoustic stimuli, suggesting a potential role in sensorimotor integration. Anatomical tracing reveals two morphologically distinct *th2+* networks, with neurons in Hc, Hi, and PT showing only local projections within the hypothalamus itself, and cells in the PON projecting broadly to a variety of CNS regions, including a number of premotor targets. We provide evidence that the motor functions of these cells are mediated by interactions between the PON and spinal projection neurons (SPNs) in the mid- and hindbrain, including neurons in the nucleus of the medial longitudinal fasciculus (nMLF), which are rapidly recruited upon activation of their dopaminergic inputs. These results elucidate a novel and essential circuit mechanism by which hypothalamic DA neurons shape vertebrate behavior.

## Results

### Phasic optogenetic activation of *th2*+ neurons evokes diverse locomotor behaviors

While our previous work demonstrated a role for the *th2*+ neurons in swim initiation, in that study we did not characterize the kinematics of the evoked behaviors in detail[19]. To better understand the types of behavior modulated by these cells we performed high-speed imaging of tail movements evoked by optogenetic stimulation. We subjected tethered larvae expressing channelrhodopsin-2 (ChR2) in *th2+* neurons (Tg[*th2:gal4; uas:chr2-eyfp*]; Fig. 1a) to 100 ms pulses of 470 nm light at intensities ranging from 0.1 to 1 mW/mm^2^ (Fig. 1b). Eyes were surgically removed 24 hours prior to the experiment to avoid the possibility of visually mediated artifacts.

**Figure 1:**
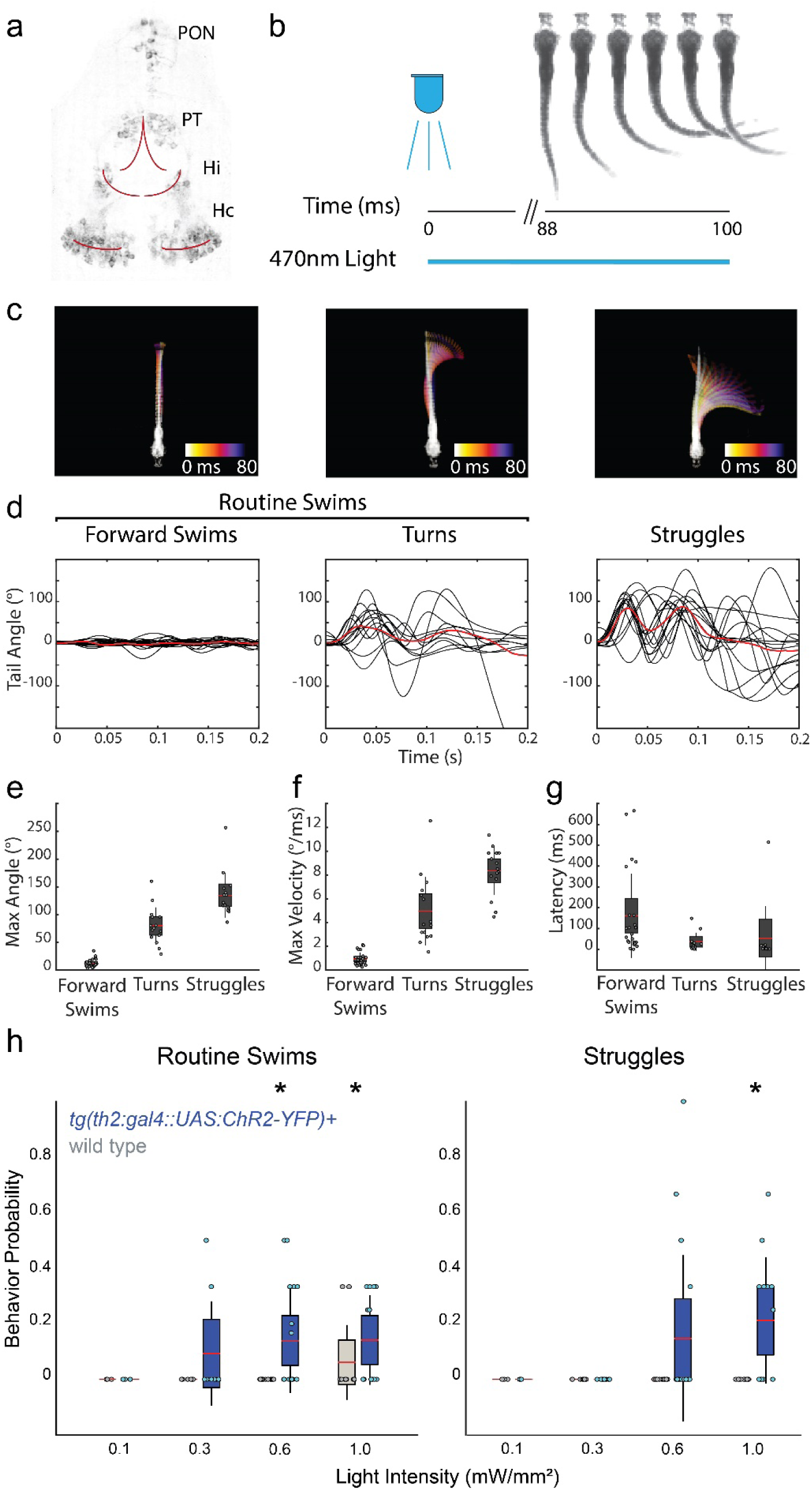
Stimulation of *th2*+ neurons drives locomotor behavior. **(a)** Two-photon fluorescence image of transgene expression in a Tg*(th2:gfp-aequorin)* animal. Expression is found in four anatomical clusters: the preoptic nucleus (PON), posterior tuberculum (PT), intermediate hypothalamus (Hi), and caudal hypothalamus (Hc). Red lines indicate approximate position of ventricles. **(b)** Experimental schematic. Eyeless fish were exposed to 100 ms pulses of 470 nm light at varying intensities, while the tail was imaged at 500 Hz. **(c)** Example behavior types evoked by stimulation of *th2*+ neurons and automatically classified by SOM clustering of kinematic parameters. Maximum intensity projections over the first 80 milliseconds, color represents time. **(d)** Tail angle traces of forward swims, turns, and struggle behaviors from all fish (n = 12 fish). Red trace, mean. **(e-g)** Max angle, max velocity, and latency of forward swims, turns, and struggles. Dots represent single trials; red bar, mean; grey boxes, 95% confidence intervals; grey lines, standard deviation. **(h)** Probability of evoked routine swims and c-bends following optogenetic stimulation in fish expressing *chr2-yfp* under the *th2* promoter as well as wild-type sibling controls. Bonferonni-corrected Kruskal-Wallis test, n = 12 fish, p < 0.01; dots represent single animals; red bar, mean; boxes, 95% CI; grey lines, SD.

We used a self-organizing map (SOM) neural network to cluster similar behaviors that we identified as forward swims, turns, and struggles based on kinematic parameters of behaviors previously characterized in freely swimming fish[24–26] (Fig. 1c, d; S1, Movie S1). Forward swims consist of a regular sinusoidal traveling wave propagating down the tail with small maximum bend angles and low angular velocity. Turns consist of a larger bend in one direction, often followed by a short bout of forward swimming. Struggles are high-velocity, high-amplitude, bidirectional movements that are sustained for hundreds of milliseconds and initiated at the caudal end of the tail, distinguishing them from other high-amplitude behaviors like c-bends (Fig. 1e, f; S1b, c).

Simultaneous activation of the entire set of *th2+* neurons evoked behaviors on relatively fast timescales, with an average latency to movement initiation of 146.7 ms (Fig. 1g). We found stimulation at 0.6 mW/mm^2^ was sufficient to evoke all of these behaviors, although forward swims and turns (which we collectively refer to as “routine swims”) were occasionally observed at lower intensities (Fig. 1h). These results suggest that *th2*+ DA neurons are broadly involved in locomotion and show that their activation is sufficient to produce routine forward swims and turns as well as higher-amplitude defensive behavior.

### Activity in *th2*+ neurons is correlated with the onset of spontaneous locomotion

We next asked if *th2*+ neuron activation is correlated with naturally occurring, spontaneous behaviors by using *in vivo* two-photon calcium imaging (Fig. 2a). Larvae expressing GCaMP5 in *th2*+ cells (Tg[*th2:gcamp5*]) were embedded in agarose with the tail freed to allow simultaneous high speed (500 Hz) measurement of behavior during the experiment. We recorded 18 minutes of spontaneous activity in *th2*+ neurons at three z-planes in each fish in order to sample from all four anatomical nuclei (Hc, Hi, PT, and PON). While frequent calcium signals were observed in Hc, Hi, and PT, spontaneous activity was infrequent and of very low amplitude in the PON (Fig. S2). Patch clamp electrophysiology of these neurons confirmed that while spontaneous activation did occur, and preceded bout initiation, it entailed a very low number of spikes per burst (Fig. S2), suggesting that a more sensitive indicator might be needed to avoid false negatives in this brain region. To address this, we transiently expressed a plasmid encoding the jGCaMP7s indicator[27] under *th2* enhancer control (*th2:jgcamp7s*) and imaged the PON in separate experimental trials. As expected, the new reporter revealed stronger and more numerous calcium transients in the PON.

**Figure 2:**
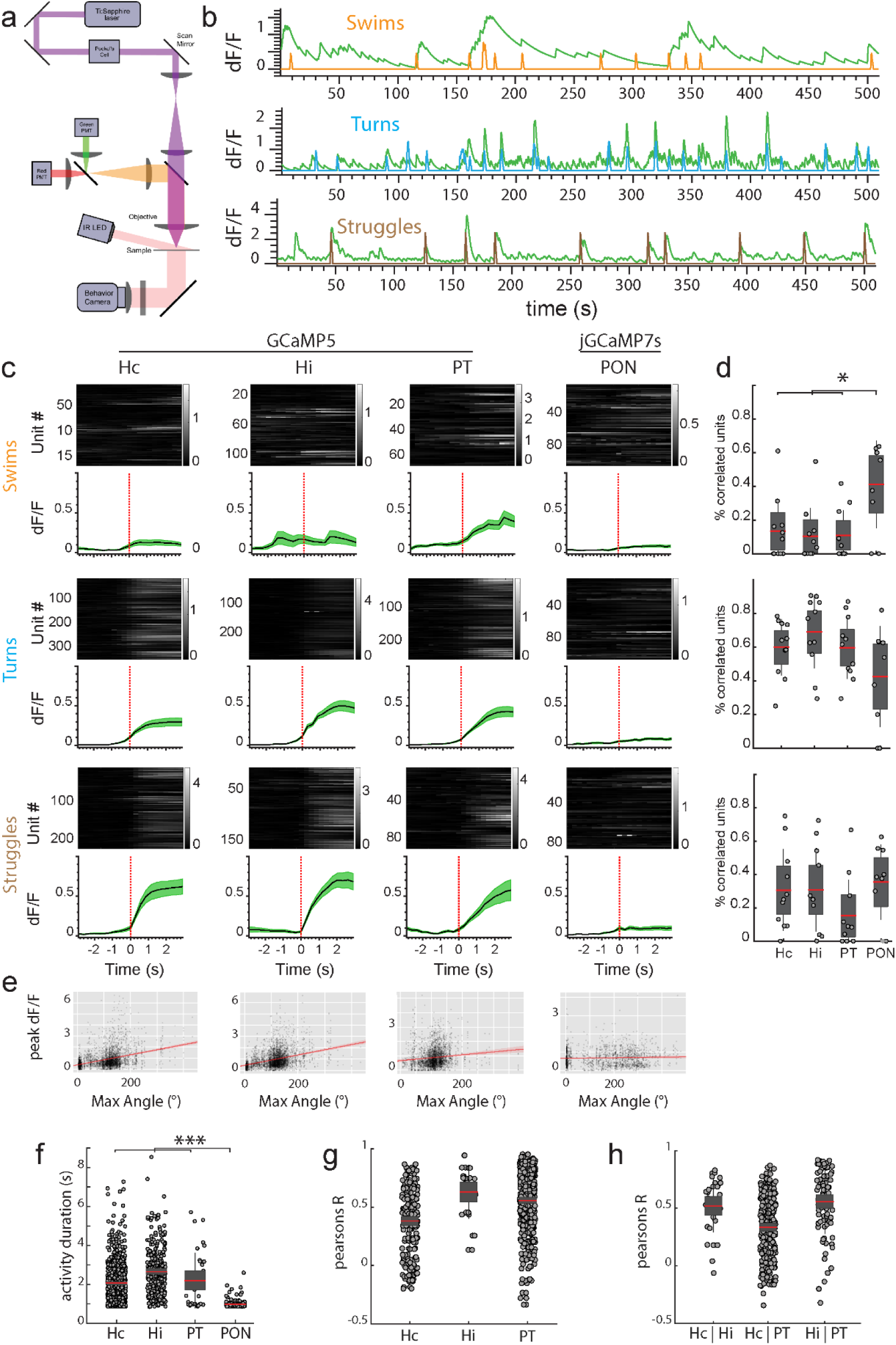
*th2*+ neuron activity is highly correlated with spontaneous behavior. **(a)** Experimental setup. Behavior recording was accomplished using a ring of oblique 830 nm LEDs mounted to the fluorescence objective. Fish behavior was recorded with a high-speed camera below the stage at 500 Hz. **(b)** Examples of forward swim, turn, and struggle regressors (orange, blue, and brown) and significantly correlated calcium traces (green). **(c)** Calcium traces aligned to spontaneous behavior onset, separated by anatomical nucleus and behavior type. Gray traces show the behavior-triggered average for individual units (mean dF/F). Green traces are mean +/-SEM across animals (n = 20 fish). Activity in Hc, Hi, and PT neurons was recorded using Tg(*th2:gcamp5*), activity in PON was recorded using transient expressors of *th2:jgcamp7s*. **(d)** Percentage of units in each nucleus significantly correlated with swim (top), turn (middle), and struggle (bottom) regressors. Red bar indicates the mean; grey boxes, 95% CI; grey lines, SD; *p < 0.05, Bonferonni-corrected ANOVA (n = 20 fish). **(e)** Scatterplot of the maximum tail bend angle of individual bouts vs the peak calcium signal of *th2*+ neurons in each nucleus within 1 second of the behavior. Red line, fitted regression. From left to right: n = 3833, 3883, 3152, & 1797 behaviors; Pearson’s R = 0.38, 0.35, 0.12, 0.05. **(f)** Average length of significant calcium transients from cells in each nucleus of Tg*(th2:gcamp5)* animals. Bonferonni corrected ANOVA, n = 1154 cells, p < 0.001. **(g)** Pearson’s R values between units within individual nuclei. Dots represent single neuron pairs. n = 200, 28, 414 pairs **(h)** Pearson’s R values between units in different nuclei. Dots represent single neuron pairs. n = 31, 263, 89 pairs

By applying the same clustering process described above, we found that fish spontaneously performed forward swims, low amplitude turns, and struggles during calcium imaging. Since qualitative observation suggested an apparent tendency for many cells to become activated during these behaviors (Movie S2), we used a regression approach to identify *th2*+ neurons with swim, turn, or struggle-associated activities. We built behavior-specific regressors by convolving the GCaMP5 impulse kernel with binary vectors equal to one at the onset of a given behavior type, and identified neurons with activity showing significantly positive correlation coefficients when regressed against these model behavior-encoding traces[28–33] (Fig. 2b). We then calculated each neuron’s behavior-triggered activity component by averaging the calcium traces surrounding spontaneous behavior onset, revealing subpopulations within each cluster that fired in concert with – often preceding – locomotor initiation (Fig. 2c).

This analysis revealed substantial differences in the nature of behaviorally-linked activity in PON versus the other three groups. In Hi, Hc, and PT there were relatively few neurons with activities associated with forward swims. (Fig. 2d), while many cells in these nuclei fired during turning and struggle behaviors. We found a positive relationship between maximum tail angle and the magnitude of the associated calcium signal in all three of these clusters (Fig. 2e). Calcium transients in neurons in Hc, Hi, and PT were significantly longer lasting than those of the PON (Fig. 2f; note that all temporal comparisons across groups were derived from Tg[*th2:gcamp5*] animals to prevent confounds arising from differences in indicator kinetics). Furthermore, activation of these cells tended to be highly synchronous, with almost all neurons in Hc, Hi, and PT showing significantly correlated activity both within and between nuclei (Fig. 2g, h; Movie S2). While subsets of neurons in the PON showed activity significantly associated with all behaviors, the PON contained a larger percentage of forward swim-associated neurons than the three more caudal clusters (Fig. 2d) and activity in that nucleus lacked a positive correlation between tail angle and the magnitude of the associated calcium burst (Fig. 2e).

Together, these results further support the idea that *th2*+ DA neurons are involved in the natural production of routine forward swimming, turning, and defensive behaviors, and also suggest that the magnitude of locomotor output is encoded in the firing rates of the three CSF-contacting clusters. Temporal correlation within and between these three groups suggest that they form a distinct functional module. Conversely, while activity in all four nuclei was phasic and associated with locomotion on fast timescales, activity in the PON did not exhibit a direct correlation with tail bend angle, suggesting that these neurons are physiologically distinct from the other three groups.

### *th2*+ neurons are activated by sensory stimuli and reduce audiomotor startle threshold

Some DAergic neurons in zebrafish respond to sensory stimuli, including *th1*+ neurons in the hypothalamus[11, 34]. To determine if the putative locomotor functions of any of the four *th2+* nuclei might be affected by sensory experience, we delivered startling acoustic/vibrational stimuli while performing two-photon calcium imaging. The intensity of the stimulus was calibrated such that it failed to evoke a behavior in approximately 50% of trials in a test fish which was not included in the calcium imaging analysis. This allowed us to identify neurons which respond to acoustic stimuli without the confound of a behavioral response. Additionally, because these measurements were performed in the same animals used for spontaneous behavior recordings, we were able to determine whether individual cells encode sensory cues, motor output, or both. Although the response rates during recording were variable, a subset of fish (n = 8) responded to approximately 50% of stimuli and were included in this analysis. Similar to the calcium responses observed during spontaneous behavior, evoked transients in Hc, Hi, and PT were of larger amplitude than those in the PON. (Fig. 3a, b). Regardless of whether a behavior was evoked, a subset of *th2*+ neurons in all four anatomical clusters appeared to consistently respond to the acoustic stimulus (Fig. 3a, b). Regression analysis quantitatively confirmed this observation (Fig. 3a, b, right panels). In combination with the results presented in Fig. 2, this suggests that the *th2*-expressing nuclei encode both sensory and motor information.

**Figure 3:**
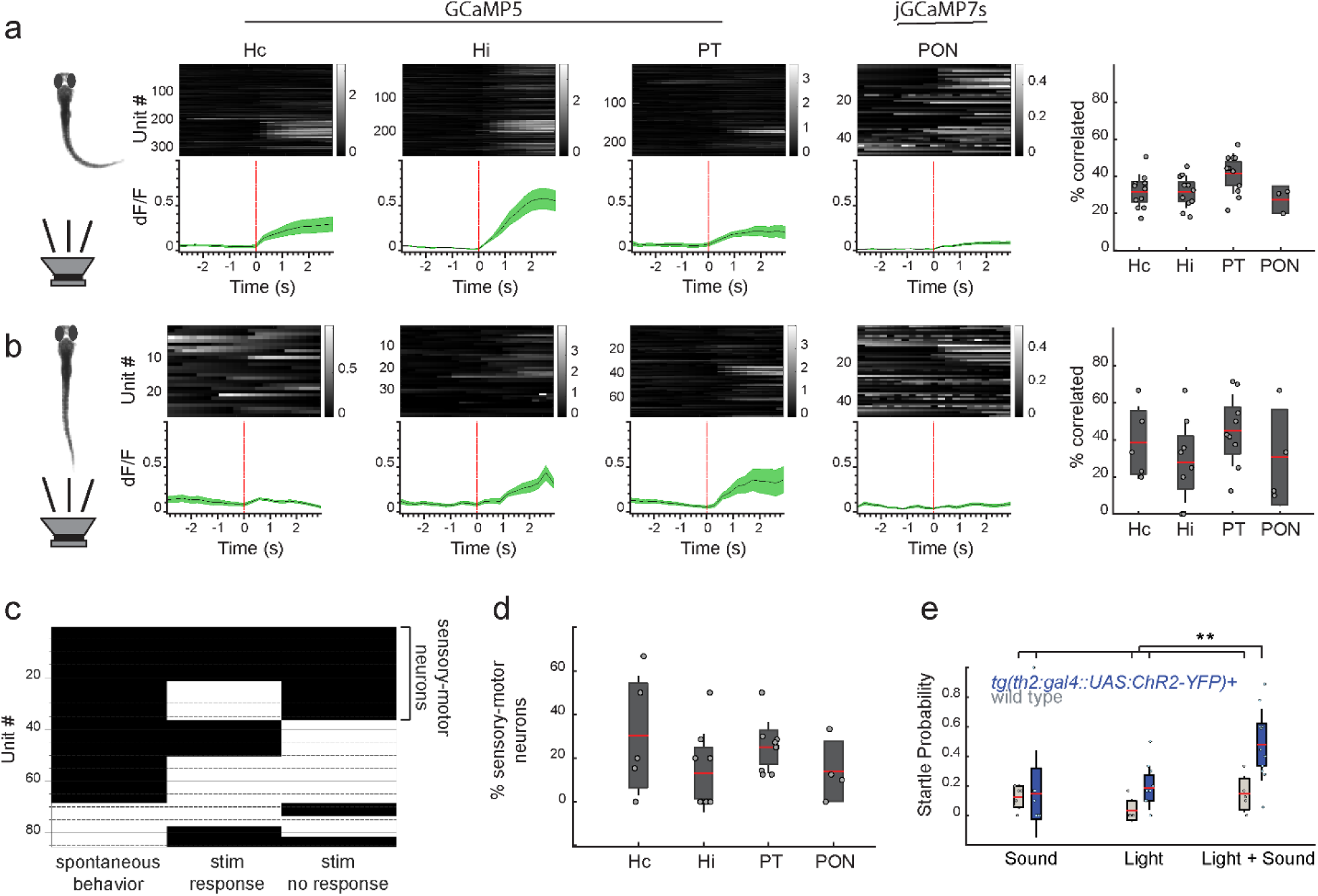
*th2+* neurons dynamically modulate audiomotor sensitivity. **(a)** Calcium traces aligned to audiomotor stimuli that resulted in a behavioral response, separated by anatomical nucleus. Gray traces show the behavior-triggered average for individual units (mean dF/F). Green traces are mean +/-SEM. Right, percent of each cluster significantly correlated with regressors encoding stimuli evoking a behavior. Pearson’s regression, alpha = 0.05. Dots represent single fish; red bar, mean; grey boxes, 95% confidence intervals; grey lines, standard deviation. From left to right, n = 11, 11, 11, 3 fish. **(b)** Calcium traces aligned to audiomotor stimuli that failed to evoke a behavioral response, separated by anatomical nucleus. Gray traces show the behavior-triggered average for individual units (mean dF/F). Green traces are mean +/-SEM. Right, percent of each cluster significantly correlated with regressors encoding stimuli not evoking a behavior. Pearson’s regression, alpha = 0.05. Dots represent single fish; red bar, mean; grey boxes, 95% confidence intervals; grey lines, standard deviation. From left to right, n = 5, 8, 9, 4 fish. **(c)** Barcode showing significant (black, p < 0.05) and non-significant (white, p > 0.05) regression results for individual units. (n = 86 neurons, 8 fish) **(d)** Percent of each cluster significantly correlated to both acoustic stimulation not evoking a behavioral response and spontaneous motor behavior. Dots represent single fish; red bar, mean; grey boxes, 95% confidence intervals; grey lines, standard deviation. From left to right, n = 5, 8, 9, 4 fish. **(e)** Probability of evoked startle behavior following audiomotor and optogenetic stimulation in fish expressing *chr2-eyfp* under the *th2* promoter as well as wild-type sibling controls. n = 5 wild type, 11 *chr2*+, Bonferonni-corrected ANOVA, p < 0.005.

Of the neurons with activity significantly associated with stimuli that did not evoke behavior, many also showed significant correlations with spontaneous locomotor events (Fig. 3c). These sensory-motor neurons made up a small subset of each cluster (Fig. 3d), but their intriguing response properties suggest the ability to modulate locomotion in a manner that depends on the animal’s recent behavioral history. We hypothesized that *th2*+ neurons might contribute to the previously observed dependence of audiomotor sensitivity on DA signaling in fish and mammals[24, 35]. To test this, we asked whether optogenetically activating the *th2*+ neurons could affect behavioral sensitivity to weak acoustic/vibrational stimuli. We exposed Tg(*th2:gal4; uas:chr2-eyfp*) fish to 100 ms pulses of 470 nm light, followed immediately by a 1 kHz sine wave delivered by a speaker mounted to the imaging platform[24, 36].The optogenetic and audiomotor stimuli were calibrated such that they rarely evoked startle responses on their own. Activating *th2*+ neurons immediately before the audiomotor stimulus significantly increased response probability in fish expressing ChR2, but not non-expressing sibling controls (Fig. 3e). These ensemble optogenetic activation data do not directly implicate the sensory-motor subgroup, but they do demonstrate an additional modulatory function for the *th2+* population as a whole that is consistent with its complex, bimodal encoding properties.

### The *th2+* projectome reveals two anatomically distinct networks

Prior to this work, the anatomical targets of *th2*+ neurons had not been thoroughly investigated. To place the behavioral functions of these neurons into a circuit context, we used a sparse labeling approach to trace their projections comprehensively[15]. One-cell stage embryos were injected with a low dose of plasmid expressing GFP-aequorin under the *th2* promoter to achieve sparse labeling of the *th2*+ population, then processed for immunohistochemical labeling of total cellular ERK content and imaged by confocal microscopy. Whole-brain z-stacks were morphologically registered to the digital z-brain reference atlas using the ERK channel as a reference (Fig. 4a, S3), and presumptive targets were identified as labeled regions in the z-brain atlas containing the termini of *th2*+ projections.

**Figure 4:**
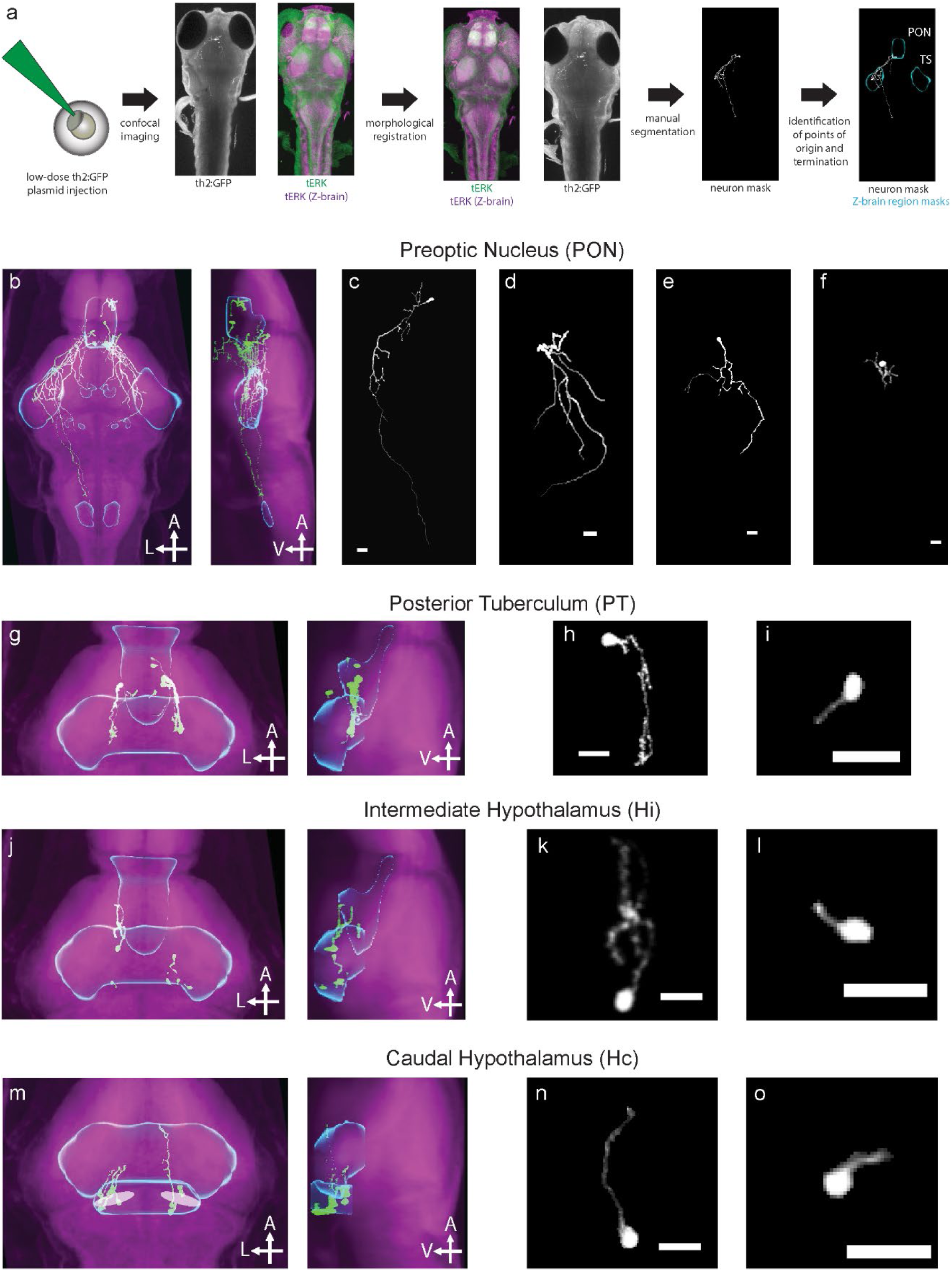
Mosaic labeling of *th2*+ neurons reveals long-range projections from the PON and reciprocal connectivity between Hc, Hi, and PT. **(a)** Experimental workflow. Fish were injected at the one cell stage with a low concentration of the *th2*:*gfp-aequorin* plasmid to achieve sparse labeling. At 6 dpf, fish were fixed and stained for GFP and total ERK. Confocal stacks were transformed for registration to the z-brain atlas using the total ERK channel and the same transformation was then applied to the GFP channel. Neurons were manually traced and the anatomical locations of the soma and axonal termini were identified using the z-brain viewer in MATLAB[38]. Cyan boundaries mark annotated brain regions from the z-brain reference atlas. **(b-e)** Maximum intensity projection of confocal stacks showing *th2*+ neurons (green) in Hc **(b)** Hi **(c)** PT **(d)** and PON **(e)** morphologically registered to the z-brain reference atlas (magenta). TS; Torus Semicircularis, nMLF; nucleus of the medial longitudinal fasciculus, IO; Inferior Olive. **(f-g)** Example ascending **(f)** and locally projecting **(g)** *th2*+ neurons in Hc. **(h-i)** Example ascending **(h)** and locally projecting **(i)** *th2*+ neurons in Hi. **(j-k)** Example descending **(j)** and locally projecting **(k)** *th2*+ neurons in PT. **(l-o)** Example *th2*+ neurons in PON. (All scale bars 20 μm)

Cells within the three CSF-contacting *th2*+ nuclei (Hc, Hi, and PT) were typified by an absence of long-range projections (Fig. 4b-d, f-k). Instead, a majority of cells had short (∼10-15 μm), local projections directed toward the third ventricle, often with a bulbous end foot contacting the ventricular space. These neurons have been previously described as liquor-contacting neurons (LCNs) of the periventricular hypothalamus[21, 37], and they were by far the most numerous cell type in this analysis (Table 1). The few projections leaving Hc were ascending and terminated in Hi. Only a single neuron was observed leaving Hi, ascending to terminate in the PT. Conversely, all neurons leaving the PT descended to terminate in Hi. These reciprocal connections might contribute to the observed correlation of activity between *th2*+ neurons across these nuclei and suggest a related function for the CSF-contacting groups.

**Table 1:** Presumptive th2+ projection targets. Points of origin are the four *th2*+ expressing hypothalamic nuclei. # denotes the total number of neurons traced from that nucleus. Points of termination are the regions of the z-brain reference atlas containing the terminal ends of *th2*+ projections. Percentages are the percentages of neurons originating from each nucleus terminating in the specified terminus region; note that single neurons often terminated in more than one region.

| Point of origin | # | Point(s) of termination | # Neurons | % Neurons |
| --- | --- | --- | --- | --- |
| Preoptic Area | 8 | Reticulospinal network: RoL1 | 2 | 25% |
|  |  | Reticulospinal network: RoM1 | 2 | 25% |
|  |  | Reticulospinal network: Nuc. MLF | 2 | 25% |
|  |  | Diencephalon | 6 | 75% |
|  |  | Preoptic Area | 2 | 25% |
|  |  | Postoptic commissure | 2 | 25% |
|  |  | Torus Semicircularis | 3 | 38% |
|  |  | Mesencephalon | 2 | 25% |
|  |  | Tegmentum | 2 | 25% |
|  |  | Migrated Posterior Tubercular Area (M2) | 1 | 13% |
|  |  | Rhombencephalon | 1 | 13% |
|  |  | Rhombomere 1, Neuropil Region 5 | 2 | 25% |
|  |  | Rhombomere 7, Neuropil Region 5 | 2 | 25% |
|  |  | Retinal Arborization field 3 | 1 | 13% |
|  |  | Eminentia Thalami | 1 | 13% |
|  |  | Tectum Striatum Periventriculare | 1 | 13% |
|  |  | Inferior olive | 1 | 13% |
| Posterior Tuberculum | 8 | Posterior Tuberculum | 4 | 50% |
|  |  | Intermediate Hypothalamus | 5 | 63% |
| Intermediate Hypothalamus | 8 | Intermediate Hypothalamus | 6 | 75% |
|  |  | Posterior tuberculum | 2 | 25% |
| Caudal Hypothalamus | 17 | Caudal Hypothalamus | 12 | 71% |
|  |  | Posterior Tuberculum | 1 | 6% |
|  |  | Intermediate Hypothalamus | 5 | 29% |

In contrast to the highly localized connectivity within the three caudal nuclei, *th2*+ neurons in the PON projected widely with highly ramified processes that descended and terminated in several brain regions, including areas with known locomotor functions (Fig. 4e, l-o). We observed projections from these neurons as far caudally as the inferior olive. Terminations were found at annotated subregions in the reticulospinal network of the hindbrain (RoL1, RoM1), and the nucleus of the medial longitudinal fasciculus (nMLF) in the midbrain, areas containing spinal projection neurons (SPNs) that have well-established roles in driving forward swimming and turning behaviors. Terminations were also observed in the torus semicircularis, known for its role in audiomotor processing. No terminations were found in the other three *th2-*expressing nuclei. This mapping of *th2*+ projections is likely comprehensive; we estimate a less than 5% chance that our approach did not sample cell types to saturation (Fig. S4). This, along with the observed differences in behaviorally-linked activity patterns, suggests that the PON and the periventricular groups have distinct functions, and identifies several brain regions that might mediate the locomotor functions of the hypothalamic DA groups.

### Brain-wide functional effects of *th2*+ neuron activity

To determine the functional significance of these anatomical projections, we used a whole-brain activity mapping (MAP-mapping) approach[38] to identify brain regions showing altered activity following stimulation of *th2*+ neurons. Fish expressing ChR2 under the *th2* promoter were exposed to blue light and subsequently stained for total ERK (tERK) and phosphorylated ERK (pERK). Using this approach, we observed subsets of neurons enriched for pERK – indicative of elevated activity – in in several brain regions (Fig. 5a-e). To quantitatively identify functionally affected areas, confocal stacks of pERK and tERK were aligned to the z-brain reference atlas and the ratio pERK/tERK was used as a normalized activity index[38] (Fig. 5f, g). This allowed voxelwise comparison of activity signals between animals, and identified several brain regions that were significantly activated across six experimental replicates.

**Figure 5:**
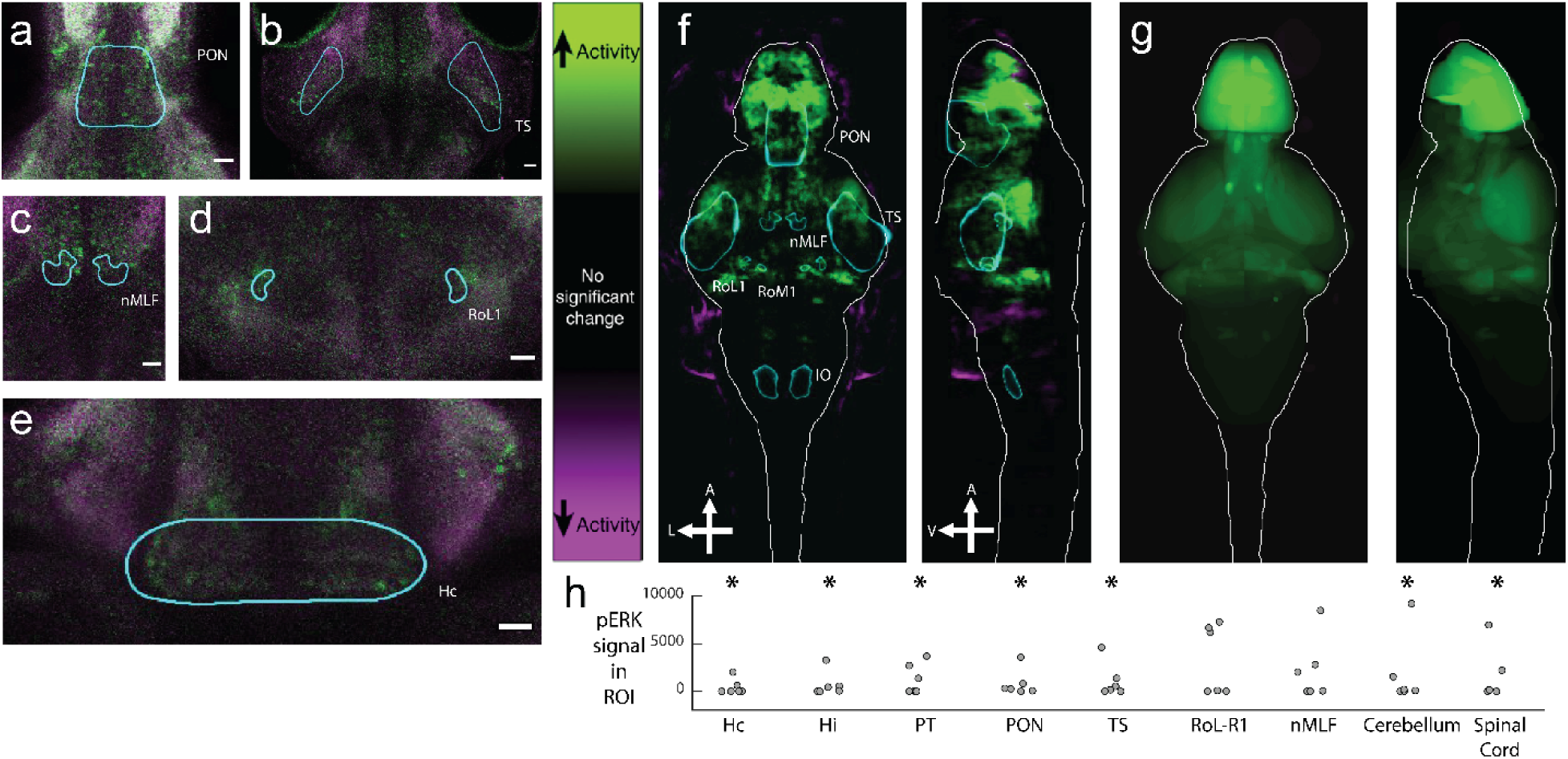
Whole-brain activity mapping confirms the functional significance of anatomically-defined target regions. **(a)** z-section showing activated neurons in the PON following optogenetic stimulation in a Tg(*th2:gal4; uas:chr2-eyfp*)+ fish. Magenta: tERK, green: pERK. **(b)** z-section showing activated neurons in the torus semicircularis following optogenetic stimulation in a Tg(*th2:gal4; uas:chr2-eyfp*)+ fish. Magenta: tERK, green: pERK. **(c)** z-section showing activated neurons in the nMLF following optogenetic stimulation in a Tg(*th2:gal4; uas:chr2-eyfp*)+ fish. Magenta: tERK, green: pERK. **(d)** z-section showing activated neurons in RoL-R1 following optogenetic stimulation in a Tg(*th2:gal4; uas:chr2-eyfp*)+ fish. Magenta: tERK, green: pERK. **(e)** z-section showing activated neurons in Hc following optogenetic stimulation in a Tg(*th2:gal4; uas:chr2-eyfp*)+ fish. Magenta: tERK, green: pERK. **(f)** Mean projection of significant, voxelwise Mann-Whitney U-statistic z-scores of an example MAP-mapping experiment. Green regions indicate increased activity following stimulation of *th2*+ neurons; purple regions indicate decreased activity (n = 15 fish per condition). **(g)** Mean projection of z-brain annotated regions showing average signal in each annotated anatomical region of the z-brain for the experiment in **(f)**. **(h)** Quantified pERK ratio in ROIs showing significantly increased pERK signal after stimulation of *th2*+ neurons. Dots are experimental replicates. Stars indicate p < 0.05, n = 6 experiments, one-tailed Wilcoxon Rank Sum Test.

Optogenetic stimulation increased activity in all four *th2+* nuclei, confirming the efficacy of the stimulus (Fig. 5h). We also observed increased activity in 69% of presumptive anatomical targets of the *th2*+ neurons, including the torus semicircularis, tegmentum, eminentia thalami, and the tectum striatum periventriculare (Table S1). The SPNs RoL-R1 and RoM1 and the nMLF were significantly activated in a subset of experiments. Other activated areas include several brain regions involved in locomotion, including the spinal cord and cerebellum. These results suggest that a majority of the anatomical projections we found act to positively modulate the activity of presumptive anatomical targets and provide a whole-brain perspective of the role these neurons play in regulating neuronal activity.

### *th2*+ afferents acutely activate spinal projection neurons

The nMLF and reticulospinal array are key mediators of the majority of locomotor behaviors, including routine swims, C-starts and other large-amplitude events[39–41]. Because our projection mapping and MAP-mapping experiments identified a potential interaction between dopaminergic neurons in the PON and targets in the nMLF, RoL1, and RoM1, we attempted to determine if these cells are dynamically modulated by *th2+* neuron activity to promote locomotion. Two-photon imaging of transgenic fish (Tg[*th2:gfp-aequorin*]) that had been backfilled with a fluorescent dextran to label the SPNs revealed numerous *th2*+ projections in close apposition to the nMLF dendrites and other SPN targets, confirming the sparse expression result (Fig. S5). Laser ablation of the *th2+* neurons in the PON reduced the frequency of spontaneous swimming, partially recapitulating our previous chemogenetic ablation result[19] and confirming the importance of these neurons for behavior initiation (Fig. S6). To more precisely determine how *th2+* afferents affect activity in these cells, we used calcium imaging and patch clamp electrophysiology to monitor changes in nMLF activity elicited by optogenetic activation of the dopaminergic inputs. Photoactivation resulted in short-latency recruitment of a large number of the spinal projection neurons, including the Mauthner cell, its segmental homologs, RoM2/3, and the nMLF (Fig. 6a-c). Calcium signals in these presumptive targets preceded simultaneously recorded fictive locomotion (Fig. 6c). We recorded the membrane voltage of MeLr, a large neuron in the nMLF that plays a well-established role in forward swimming [39, 40], in a current clamp configuration during optogenetic stimulation of *th2*+ neurons. Stimulation evoked sustained bursts of spiking in MeLr reliably within 100 ms of stimulation onset (Mean, 31.68 +/-8.19 ms; Fig. 6d-g). These results suggest that the acute locomotor functions of *th2*+ DA neurons are mediated at least in part by a projection from the preoptic nucleus of the hypothalamus to the nucleus of the medial longitudinal fasciculus and other spinal projection neurons.

**Figure 6.**
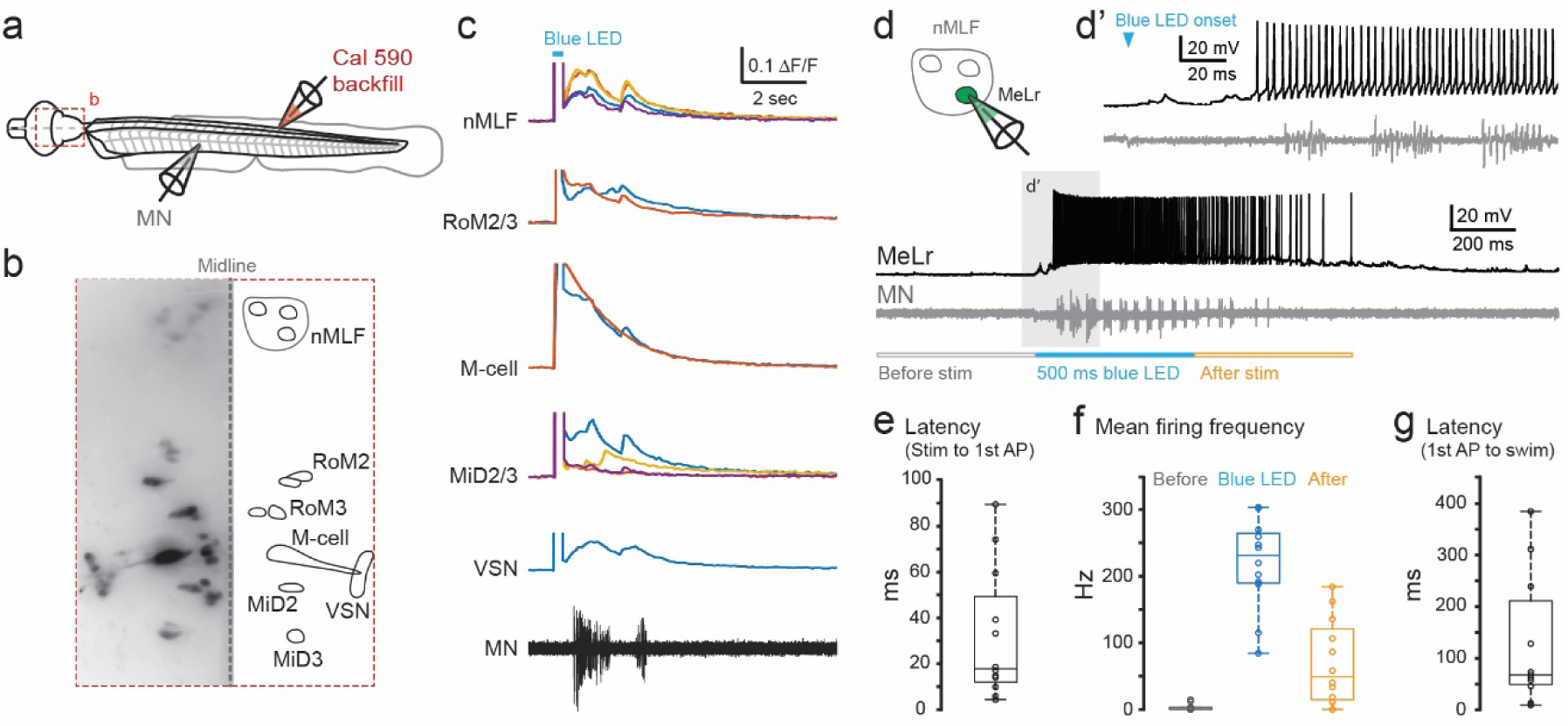
*th2*-expressing dopamine neurons drive locomotion by activating spinal projection neurons in the mid- and hindbrain. **(a)** Schematics of the experimental preparation for simultaneous calcium imaging of spinal projection neurons (SPNs) and motor nerve (MN) recordings. SPNs are labeled by backfilling from the spinal cord with the dextran conjugated calcium sensitive dye Cal-590. The red box highlights the brain region expanded in (b). **(b)** Left: maximum intensity projection image of SPNs in the midbrain and hindbrain. Right: cell bodies of imaged SPNs are outlined. VSN: vestibulospinal neuron. **(c)** Normalized calcium fluorescence change versus time for identified SPNs shown in (b) during optogenetic stimulation in Tg(*th2:gal4; uas:chr2-eyfp*) fish. Uniquely colored traces indicate different neurons within each class. The bottom black trace represents the motor nerve recording; the blue box indicates the photoactivating stimulus, also apparent in the artifactual increase in fluorescence signal for each cell. **(d)** Whole-cell current clamp recording from a representative nMLF neuron, MeLr, during optogenetic stimulation of *th2*-expressing neurons with simultaneous motor nerve recording. (d’) An expanded view of the shaded area in the lower panel. **(e)** Box plot showing the latency from the photoactivating stimulus onset to the first MeLr action potential. n = 4 fish, 12 trials. **(f)** Box plot showing mean firing frequencies of the MeLr in the 500 ms time windows before, during, and immediately after photoactivating stimulation. n = 4 fish, 12 trials. **(g)** Box plot showing the latency from the first MeLr action potential to the onset of motor nerve activity. n = 4 fish, 11 trials.

## Discussion

The dopaminergic cell groups of the vertebrate hypothalamus are poorly understood with respect to their function and physiology. Our previous work showed that the subset of these neurons defined by *th2* expression in zebrafish are required to maintain a normal frequency of spontaneous locomotor initiation. Here, we show that *th2+* cells are involved in the production of a range of swimming behaviors and participate in sensorimotor processing. While neurons in all four *th2*-expressing hypothalamic nuclei fire during spontaneous and evoked swimming, differences in their activity and connectivity suggest that these cells fall into two functionally distinct groups. Neurons in Hc, Hi, and PT comprise an interconnected, periventricular network that appears to contribute to locomotion by unknown means, while cells in the preoptic nucleus modulate the probability of swim initiation through projections to the nucleus of the medial longitudinal fasciculus (nMLF) and the reticulospinal array. We find that locomotor control by these cells is fast; phasic activity begins within a few hundred milliseconds of behavior onset and is sufficient to drive spiking in the nMLF and swimming behavior within a similar timeframe. Our findings are the first to delineate a specific role for these cells in sensorimotor behavior and posit a novel mechanism by which DA neurons in the vertebrate PON shape the activity of a premotor network.

### *th2*+ neurons are involved in the production of behavior on fast timescales

Neuromodulators like DA are generally presumed to act on longer timescales than “classical” neurotransmitters due to the nature of metabotropic transmission and the often slow, tonic activity patterns of modulatory neurons, which ostensibly affect behavior over the course of many seconds, minutes or hours[42]. However, these cells are also capable of firing phasic bursts of activity[22, 43, 44], which have recently been shown to exert acute effects on locomotor function in the rodent striatum[22, 44] and the zebrafish spinal cord[5, 11]. Our finding that many of the *th2+* neurons show bursts of phasic activity that are well correlated with spontaneous and evoked behavior at latencies of tens to hundreds of milliseconds, often preceding the initiation of movement, suggest that such dynamics might represent a general mechanism by which DAergic neurons promote locomotion.

Optogenetic manipulations further corroborate the relevance of phasic DA activity to activity in downstream circuits and locomotor behavior. The typical latency between the onset of the ChR2 stimulus to activation of the nMLF ranged between 20 – 100 ms, fast enough to account for the observed behavioral latency of around 140 ms. These values are consistent with a modulatory mechanism, though the true, physiological latencies could potentially be masked somewhat by an additional lag in ChR2-evoked spiking in the *th2+* neurons themselves [45]. Conversely, while the effects seen here are significantly slower than the sub-10 ms timescales typical of “classical” neurotransmission, the specific contribution of DA versus e.g. a fast-acting co-transmitter such as glutamate remains to be seen. It will be also be important to explore the persistence of DA-mediated sensitization of the motor circuit in the future. Given uncertainty surrounding the temporal features of volumetric neurotransmission in most cases[42], working within this established circuit should allow us to gain insight into a fundamental parameter of DA signaling. Additionally, while we focused our investigation on discrete locomotor bouts, it remains possible that the cumulative effect of DA released during successive, phasic bursts, or perhaps an as-yet unidentified slower modulation of “tonic” firing rates in these neurons, affects other behavioral functions that have been attributed to these regions of the hypothalamus, such as wakefulness or hunger[46, 47].

Our data also support an acute, phasic role for the *th2+* neurons in audiomotor modulation, and identify a novel neuroanatomical source for the DA signals that have long been known to influence sensorimotor behaviors in fish[24, 48]. The ability of a subset of individual *th2*+ neurons to encode both sensory and motor information suggests that these cells might be capable of dynamically integrating sensory information and modulating motor activity in response to a changing behavioral context, although a conclusive demonstration of this hypothesis awaits targeted manipulation of the sensory-motor cell group. The diverse anatomical projections noted for cells in PO, along with the complexity of coding properties in this group, imply a correspondingly high degree of functional heterogeneity that remains to be explored.

### *th2*+ cells in PT, Hi, and Hc comprise an interconnected, CSF-contacting network

We find a striking degree of reciprocal connectivity between neurons in the CSF-contacting nuclei PT, Hi, and Hc via projections that likely correspond to the endohypothalamic tract[49, 50], but do not observe direct connections with other brain regions. In contrast to the *th1*+ neurons of PT, no descending spinal projections from the tubercular *th2+* cells were observed[15], an unexpected difference between genetically defined subgroups of DAergic neurons within a single anatomical nucleus. Our observation that many *th2*+ neurons in the periventricular regions fire during swim initiation is therefore surprising because it suggests an acute behavioral role for neurons that do not project outside of the hypothalamus. This is consistent with previous observations of other locally-projecting hypothalamic DA neurons with locomotion-associated activity[11]. However, the lack of a distal target with a known locomotor function leaves the means by which these cells might control swimming unresolved.

One potential clue lies in the overwhelming prevalence of cells with short, local projections into the ventricle within all three of these nuclei. These CSF-contacting DAergic neurons have been described previously[21, 37] although until the discovery of the *th2* gene[16, 17, 20] they were thought to accumulate DA from the CSF rather than produce it autonomously. One possibility is that these neurons modulate the activity of local neurons which themselves project outside the hypothalamus to affect behavior[51]. However, an intriguing alternative that is consistent with our data is the possibility that these cells influence locomotion using a completely novel functional mechanism in which DA is secreted into the ventricle and signals by diffusion through the CSF. It is unclear what the functional targets for such a mechanism might be, or whether any candidate structures lie close enough to the 3^rd^ ventricle for a diffusion-based mechanism to have acute behavioral effects. However, recent observations of CSF transport dynamics in larval zebrafish have shown that such processes can occur far more rapidly than previously appreciated[52].

### A functionally distinct population of preoptic *th2*+ neurons drives behavior via projections to the mid- and hindbrain

Here we find that *th2*+ DA neurons in the preoptic region of the hypothalamus are critically involved in the initiation of locomotor behavior. A number of observations in mammals suggest that this behavioral role could be evolutionarily conserved. While previous anatomical characterization of the mammalian preoptic equivalents in DAergic groups A14/A15 has been somewhat anecdotal and incomplete, it is known that these cells also project to a variety of brain regions, including the pituitary[14], thalamus[53], habenula[54], amygdala, and nucleus accumbens[55]. Except for the pituitary projections, which are the primary source of DA released into the adenohypophysis to regulate lactation, the functional significance of these projections is unclear. Electrical stimulation of the preoptic area elicits locomotor behavior in the rat[6], but no functional studies had explored how the DA neurons in that area might contribute. Our data suggest that these cells play an essential role in motor initiation and should motivate further study of the role of preoptic sources of DA in locomotion in other vertebrates.

While we chose to focus on interactions with the nMLF and reticulospinal array, other presumptive targets identified in our anatomical and functional mapping experiments are likely relevant for the behaviors we observe during optogenetic manipulation. Particularly interesting is the projection to the torus semicircularis, a brain area known to mediate audiomotor sensory processing[56]. This area is responsive to auditory stimuli in larval zebrafish[57] and could contribute to the modulation of startle sensitivity in parallel with direct projections onto the SPNs.

Our data provide essential details that were missing from our understanding of modulatory inputs within the mid- and hindbrain. Dopaminergic innervation of the SPNs had been previously suggested by immunohistochemical staining against *th1* and some functional observations of Mauthner cell activity[48, 58, 59], though the source of modulation in those cases was unclear[60]. Prior comprehensive tracing of *th1*+ DA neurons did not employ the anatomical alignment and atlasing approach used here and so innervation of small cell clusters such as the nMLF could not be identified; in that work, *th1+* cells in the PON were observed projecting only to the postoptic commissure, endohypothalamic tract, and hypothalamus[15]. Since some *th2*+ neurons in PON also express *th1*[20], and since our single-cell imaging experiments approached saturation, it is possible that a subset of the SPN-projecting cells are marked by both tyrosine hydroxylase isoforms and account for the innervation observed previously.

The unique activity dynamics in the PON population further suggest a functional distinction from the three more caudal nuclei. While *th2*+ neurons in Hc, Hi, and PT showed high-amplitude calcium transients, calcium imaging and electrophysiology revealed relatively low-frequency, low-amplitude, locomotion-associated spiking in the PON. Additionally, while the amplitude of calcium transients in Hc, Hi, and PT was positively correlated with the vigor of behavioral events, transients in the PON were of consistent magnitude regardless of behavior type. This, along with their widely distributed projections and lack of anatomical connectivity with the three other nuclei, suggests that these neurons comprise a distinct functional group that may broadly facilitate sensorimotor processes to promote behavioral arousal on fast timescales. This is consistent with previous work showing that the spinally-projecting *th1*+ DAergic neurons in the PT are functionally distinct from nearby locally-projecting *th1*+ neurons in the dorsomedial hypothalamus (Hdm) and Hc. *th1*+ PT neurons are functionally diverse; some respond to sensory stimulation and some fire prior to spontaneous swimming. In contrast, the locally-projecting neurons in Hdm and Hc primarily fire during spontaneous locomotion[11]. Physiological and functional differences between far-projecting and locally-projecting DAergic neurons may be a general feature of DAergic neurons and deserves further investigation.

Together, these data show that *th2*+ DAergic neurons in the preoptic nucleus of the hypothalamus exert fast sensorimotor control in part via a newly-identified projection onto pre-motor neurons in the brainstem. Future work will further characterize the contribution of these interactions to specific forms of sensorimotor processing and behavior selection, and the means by which signaling at these targets is integrated with activity in the other functional efferents of the hypothalamic DA neurons.

## STAR ★ Methods

### LEAD CONTACT AND MATERIALS AVAILABILITY

Further information and requests for reagents may be directed to, and will be fulfilled by Dr. Adam Douglass. All unique/stable reagents generated in this study are available from the lead contact without restriction.

### EXPERIMENTAL MODEL DETAILS

#### Fish husbandry

Zebrafish larvae were raised in E3 media until 8 days post fertilization (dpf) and fed with rotifer co-culture from 5-8 dpf. Larvae and adult fish were maintained on a 14-hour light/ 10-hour dark cycle at 28 ⁰C. All experiments were performed on 6-8 dpf larvae with the exception of electrophysiological experiments, which were performed on 4-5 dpf larvae. All protocols and procedures involving zebrafish were approved by the University of Utah Institutional Animal Care and Use Committee.

#### Transgenic lines and plasmids

The transgenic lines Tg(*th2:gfp-aequorin*)^zd201^ and Tg(*th2:gal4-vp16*)^zd202^, along with the corresponding plasmids, are described previously[19]. To create the *th2:jgcamp7s* plasmid, the coding sequence was amplified by PCR to introduce flanking SpeI and SacII sites, then digested and ligated into the corresponding sites of *pDest-Tol2*. The resulting destination vector was then LR-reacted with *pENTR:th2(9.6kb)*[19] using a Gateway cloning kit (Thermo Fisher). *th2:gcamp5* was created by the LR reaction of *pENTR:th2(9.6kb)* with *pDest-Tol2:gcamp5*[61]. The Tg*(th2:gcamp5)* transgenic line was created by co-injecting 30 pg of plasmid with 30 pg of Tol2 RNA into single-cell embryos, identifying adult founders by screening for fluorescence in their offspring, and outcrossing for two subsequent generations. For transient expression experiments, *th2:gfp-aequorin* or *th2:jgcamp7s* were injected at 1 ng/μL or 30 ng/μL, respectively, and larvae expressing the transgene in an appropriate number of neurons were identified by screening at 3-5 dpf.

### METHOD DETAILS

#### Optogenetic manipulation of behavior

Transgenic or wild-type larvae were mounted in a drop of 1.5% low-melt agarose (LMA) and the tail was subsequently freed to allow monitoring of tethered behavior. Eyes were surgically removed 24 hrs prior to experimentation to avoid confounds arising from direct visual stimulation by the optogenetic impulse. Fish were imaged at 500 fps using a Pike F032B camera (Allied Vision Technologies). Optogenetic stimuli were delivered using a 470 nm PCB-mounted LED (Luxeon) and calibrated using a ThorLabs S310C power meter. To evaluate startle modulation, acoustic/vibrational stimuli were delivered via a speaker mounted directly to the imaging platform. All stimulus waveforms were generated using a National Instruments PCIe-6323 data acquisition board and custom LabView software that synchronously controlled the camera acquisition.

#### *In vivo* two-photon calcium imaging

Transgenic animals (*Tg[th2:gcamp5]* or transient expressers of *th2:jgcamp7s*) were immobilized in 1.5% low-melting point agarose and the tail subsequently freed to allow for simultaneous recording of behavior. Calcium fluorescence was recorded using a custom-built two-photon microscope at 3.44 Hz. Fluorescence was recorded at each of three z-planes in order to sample from each of the four anatomical *th2*+ nuclei. One plane included Hc and Hi, the second plane included PT, and the third plane included PON. We recorded 17 minutes of spontaneous activity at each plane. Acoustic stimulation experiments were performed concurrently in the same fish. These consisted of nine 10ms, 1 kHz sine waves delivered through a speaker mounted directly to the imaging platform at each z-plane. The intensity of these stimuli were calibrated such that they evoked behavior in 50% of trials in a test fish which was not included in the calcium imaging analysis. For analysis of stimulus-evoked activity during trials evoking a behavioral response or failing to evoke a behavioral response, we considered for analysis only neurons which were recorded during at least three stimuli evoking a response or failing to evoke a response. For analysis of sensory-motor neurons, we included only neurons recorded during at least 3 trials which successfully evoked a behavior and at least 3 trials which failed to evoke a behavior. Activity sources were automatically detected and traces were extracted using the CaImAn package in MATLAB[62]. The accuracy of CaImAn-extracted dF/F traces was verified manually in a subset of trials using manual ROI selection and a simplified dF/F calculation. Findings regarding correlation with spontaneous behavior and acoustic stimuli were reproduced with another calcium imaging analysis package[63] to ensure robustness (data not shown) and to measure the lengths of significant calcium transients. Transients were defined as significant upon meeting two conditions: (1) They cannot be explained as random fluctuations in the signal over time, and (2) Their temporal features match those of the reporter. To evaluate these conditions, we calculated a dynamic threshold that depends on both intensity variance within each ROI and the biophysics of the calcium reporter using a Bayesian odds ratio framework.

#### Behavior quantification, and classification

For behavioral recordings acquired during both calcium imaging and optogenetics experiments, tail angles were extracted and analyzed using custom MATLAB scripts. Eight evenly-spaced tracking points were selected along the center axis of the tail using a previously-described algorithm[25]. Cumulative tail angle was calculated as the sum of the angles between these tracking points. These measurements were used to extract the following kinematic parameters for each swim bout: Max velocity, mean velocity, max angle, end angle, the max angles at each individual segment 1-7, max segmental angular change, mean segmental angular change, duration. Swim bouts were then divided into half-beats and the same parameters were extracted for the first three half-beats. This resulted in a total of 59 kinematic parameters per behavior, which were used to train a self-organizing map neural network for clustering of behavior types. We found that a 3×3 output layer obtained 9 initial clusters comprising three types of behavior. Clusters with similar max angle and max velocity were merged manually. The network was trained over 20,000 iterations. Behaviors from each experiment were clustered separately.

#### Immunohistochemistry

Fish age 6 dpf were fixed in 4% paraformaldehyde (PFA) in PBS + 0.25% Triton-X (PBST) overnight at 4⁰C. Fish were then washed in PBST, permeabilized in 0.05% Trypsin-EDTA on ice for 45min, washed in PBST, blocked in PBST + 1% bovine serum albumin (BSA) + 1% DMSO + 2% normal goat serum (NGS), and then incubated overnight in primary (1:500 dilution) and secondary (1:1000 dilution) antibodies in PBST + 1% BSA + 1% DMSO. The following antibodies were used.

Anti-tERK: ZIRC antibody collection, Cell Signalling #4696.

Anti-pERK: ZIRC antibody collection, Cell Signalling #4370.

Anti-Acetylated tubulin: SigmaAldrich #T7451.

#### MAP-mapping

6 dpf larval zebrafish were fixed immediately following 30 minutes of optogenetic stimulation (1 second pulses, 5 second inter-stimulus interval) and stained according to the protocol above. Whole-mount fish were mounted in 1.5% (wt/vol) low-melting-point agarose, sealed with a glass coverslip, and imaged on an inverted confocal microscope (Leica SP8) using a 20x/1.0-NA water immersion objective. Fish were imaged at a voxel size of 0.78 x 0.78 x 2 microns (x y z). Two imaging tiles were acquired and stitched together using Nikon NIS Elements software. Pre-processing and analysis of MAP-maps were performed as described in Randlett et al[38] using the provided ImageJ and MATLAB scripts. Voxelwise Mann-Whitney U-statistic Z-scores were calculated for pairwise comparisons between the pERK/tERK ratio in ChR2-expressing and non-expressing sibling controls. Significant z-scores were identified using a bootstrapping approach with a false discovery rate threshold of 0.005%[38]. We performed six identical MAP-mapping experiments. A one-tailed Fischer’s exact test against the hypothesis of zero signal was used to identify regions of the z-brain with consistent elevations in activity across experimental replicates[64].

#### Whole-brain anatomical registration and projection tracing

Confocal stacks were aligned to the z-brain reference atlas using CMTK (http://www.nitrc.org/projects/cmtk/)[65] in an Ubuntu Linux environment. We performed non-rigid registration using the command string (-awr 0102 -X 52 -C 8 -G 80 -R 3 -A ’--accuracy 0.4’ -W ‘--accuracy 1.6’)[38]. The z-brain 6-dpf nacre anti-tERK reference stack was used as a template.

For projection tracing, early embryos were injected with *th2:gfp-aequorin* to achieve sparse labelling, then immunostained for tERK and GFP at 6 dpf, then imaged by confocal microscopy. Neurons with non-overlapping projections were selected for tracing using the simple neurite tracer plugin in FIJI[66]. Brain regions containing cell soma and projection termini were identified using the z-brain viewer in MATLAB.

#### Laser ablations

6 dpf Tg*(th2:gfp-aequorin)* larvae were immobilized in 1.5% low melting point agarose and anaesthetized using MS-222. Neurons were located by two-photon microscopy, and then ablated by repeatedly scanning a circular path around the cell periphery with the laser tuned to 920 nm at an intensity of 50 mW. For sham ablations, this procedure was repeated at an equivalent number of ROIs in the PON that did not contain labeled neurons.

#### Calcium imaging of spinal projection neurons

Spinally-projecting neurons in the hindbrain and midbrain were retrogradely labeled in Tg*(th2:gal4; uas:chr2-yfp)* zebrafish larvae at 4 dpf by injecting the red calcium dye Cal-590 (dextran conjugate 3,000 MW, AAT Bioquest; 10% w/v in patch solution without EGTA, composition in mM: 125 K-gluconate, 4 MgCl2, 10 HEPES, 4 Na2ATP, adjusted to pH 7.2 with KOH) into the spinal cord at around body segment 15. At 5-7 dpf, injected larvae were immobilized in 0.1% w/v α-bungarotoxin (Tocris) and then stabilized on a Sylgard-lined glass bottom dish containing extracellular solution (composition in mM: 134 NaCl, 2.9 KCl, 1.2 MgCl2, 2.1 CaCl2, 10 HEPES buffer, 10 glucose, adjusted to pH 7.8 with NaOH). The brain was dissected out to remove all sensory inputs from the head area with the connection between the brain and the spinal cord preserved. Calcium imaging was performed using an upright microscope (BX51WI, Olympus) with a sCMOS camera (Zyla, Andor DG-152X-C1E-Fl) and an LED light source (SOLIS-3C, THORLABS). Calcium responses of the reticulospinal neurons were acquired with Solis Software (Andor) at 10 Hz and analyzed using custom MATLAB scripts. Motor nerve activity was acquired simultaneously with calcium imaging at 50 kHz (for details see Electrophysiology Recordings). To activate TH2 neurons LuxeonStarLEDS) triggered by a digital I/O device (USB-6501, National Instrument) with custom LABVIEW scripts was used.

#### Electrophysiology

4-5 dpf Tg*(th2:gfp-aequorin)* larval zebrafish were used for TH2 neuron recordings. Zebrafish larvae were immobilized in α-bungarotoxin and placed on a Sylgard-lined glass-bottom dish containing extracellular solution. The brain was exposed and flipped ventral-side-up. For MeLr recordings, Tg*(th2:gal4; uas:chr2-yfp)* larvae were retrogradely labeled with Texas Red Dextran as described above. After mounting, the head was rotated dorsal-side-up and secured with pins, and the eyes were removed to prevent visually evoked responses.

The preparation was then transferred to the physiological recording apparatus, comprised of an upright microscope (BX51WI, Olympus) equipped with a 40x/0.8 NA water immersion objective and two motorized micromanipulators (MPC-385 and TRIO-245, Sutter Instruments). For whole-cell patch recordings, standard wall glass capillaries were pulled with a pipette puller (PIP6, HEKA) to make recording pipettes with resistances between 14-16 MΩ, which were then back-filled with patch solution (composition in mM: 125 K-gluconate, 4 MgCl_2_, 5 EGTA, 10 HEPES, 4 Na_2_ATP, adjusted to pH 7.2 with KOH). Na3GTP (final concentration 0.3 mM) and AF488 Hydrazide (20uM) were added into the patch solution immediately before each experiment. For peripheral motor nerve recordings, glass capillaries were pulled as for the patch pipettes, and an opening of approximately 30 µm was made at the tip with a beveller (EG-44, Narishige). Electrophysiological recordings were acquired at 50 kHz using a Multiclamp 700B amplifier, a Digidata 1550B digitizer, and pClamp software (Molecular Devices). Electrophysiological data were analyzed using DataView (Heitler, nm, Thorlabs) was used to activate TH2 neurons expressing ChR2.

## QUANTIFICATION AND STATISTICAL ANALYSIS

No statistical methods were used to predetermine sample sizes, but our sample sizes were based on prior literature and established best practices in the field[11, 33, 38]. Calcium traces are shown as mean +/-s.e.m., distributions are shown as mean, 95% CI, and 1 SD. Plots and statistics were generated in MATLAB and Python 3. Comparisons of multiple groups were performed using one-way or two-way ANOVA, followed by Bonferroni’s multiple comparisons test when significance was detected. Regressors for calcium imaging analysis were constructed by convolving the appropriate GCaMP impulse kernel with a binary vector encoding behavior onset frames[29]. Acoustic stimulus regressors were constructed over a six second window around the stimulus and regressed against the stimulus-triggered average of recorded units. Pearson’s regression coefficients and associated p-values were calculated using the corr function in MATLAB. We performed six MAP-mapping experimental replicates and extracted significant pERK signal in each ROI of the z-brain as described previously[38]. These six results were then tested using a one-tailed Wilcoxon signed rank test against the null hypothesis of 0 signal in each ROI, alpha = 0.05.

## DATA AND CODE AVAILABILITY

Data and custom-built MATLAB scripts are available from the authors upon request.

## Acknowledgements

This work was funded by the National Institute for Neurological Disorders and Stroke (F31-NS100412 to J.B.), the National Institute of Mental Health (R56-MH110529 to A.D.D.), and the Sloan Foundation (A.D.D.). Special thanks to J. Bonkowsky and R. Dorsky for many insightful conversations, feedback on the manuscript, and sharing of primary antibodies and other reagents.

## Author Contributions

The project was conceived by J.P.B and A.D.D. J.P.B. performed the majority of experiments and data analysis. W.W. acquired and analyzed all electrophysiology data. R.E. performed MAP-mapping stimulation experiments and assisted with laser ablation data analysis. W.W. collected SPN calcium imaging data and A.D.D. analyzed the results. E.R. cloned the *th2*:j*gcamp7s* plasmid and performed single-cell embryo injections. J.B., W.W., R.E., and A.D.D. wrote and edited the manuscript.

## Notes

### Competing Interest Statement

The authors have declared no competing interest.

