## Supplemental Figures for "Hypothalamic dopamine neurons control sensorimotor behavior by modulating brainstem premotor nuclei"

**Figure S1: Clustering of behavior types using a self-organizing map (SOM) neural network**

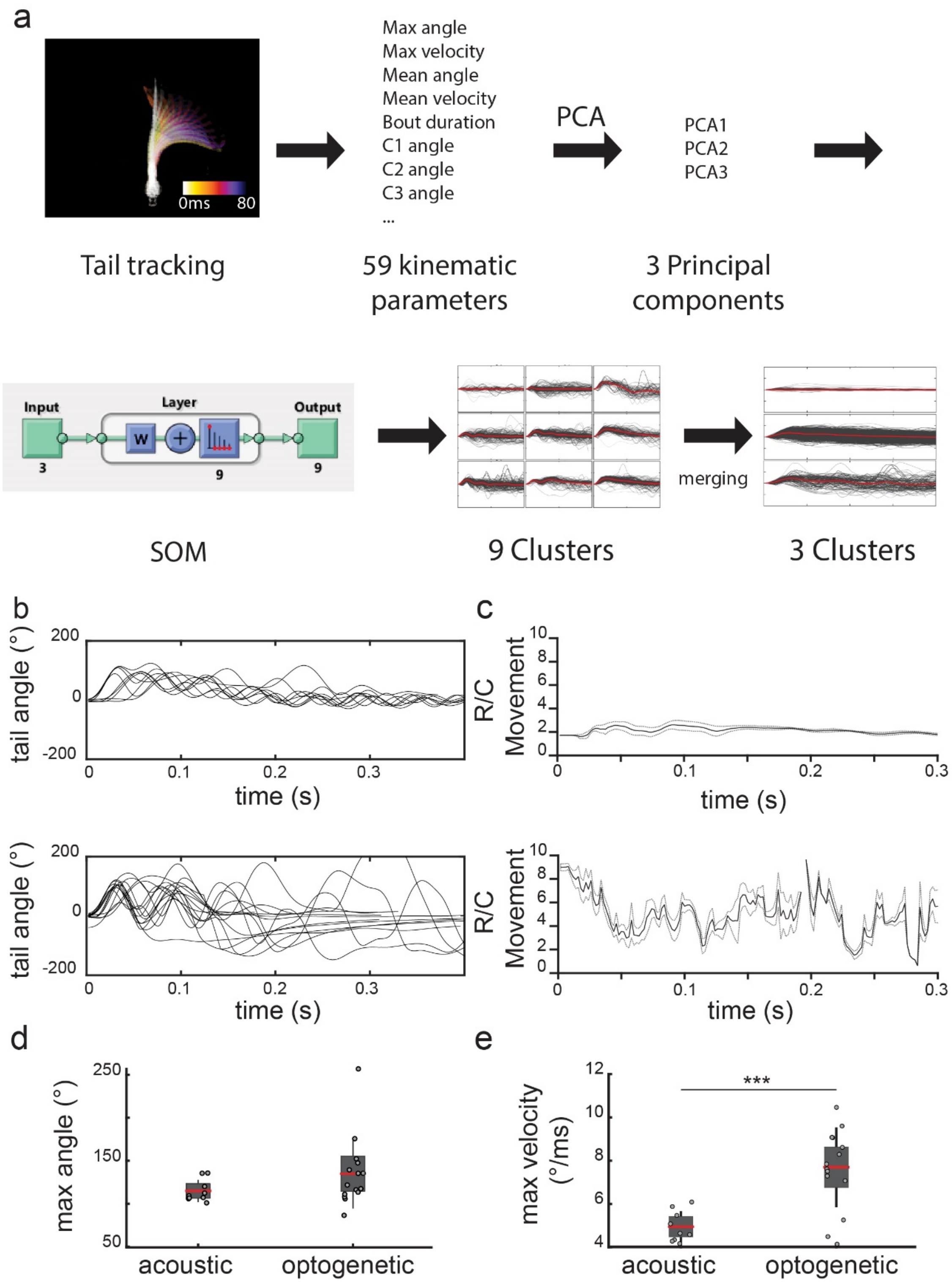

**Figure S1: Clustering of behavior types using a self-organizing map (SOM) neural network**

(a) Workflow diagram of behavior quantification and classification. Kinematic parameters were measured using 8 equally spaced tracking points along the length of the tail. The total tail angle was calculated as the cumulative sum of the angles between these tracking points. We calculated a total of 59 kinematic parameters from each swim bout and reduced the data to three principal components. These three principal components explained >90% of the variance in kinematic parameter space. We then used these PCs to train a self-organizing map neural network to separate behaviors into 9 clusters to ensure oversegmentation, and then finally reduced these 9 clusters into 3 based on visual similarity of the traces and statistical comparison of initial bend angles, maximum tail angles, and maximum velocities. Clusters were only combined if these values were statistically indistinguishable.

(b) Tail angle over time traces of acoustically evoked startle (top) and optogenetically evoked struggle behaviors (bottom).

(c) Ratio of distance moved by the caudal tip of the tail to distance moved by the rostral end of the tail for acoustically evoked startle (top) and optogenetically evoked struggle behaviors (bottom). Solid line, mean; dashed line, SEM.

(e) Maximum tail angle of startle vs. struggle behaviors ($n = 9$, $n = 15$, $p > 0.05$)

(e) Maximum tail velocity of startle vs. struggle behaviors ($n = 9$, $n = 15$, $p < 0.001$)

**Figure S2: Spontaneous, locomotion-associated activity of preoptic *th2*+ neurons**

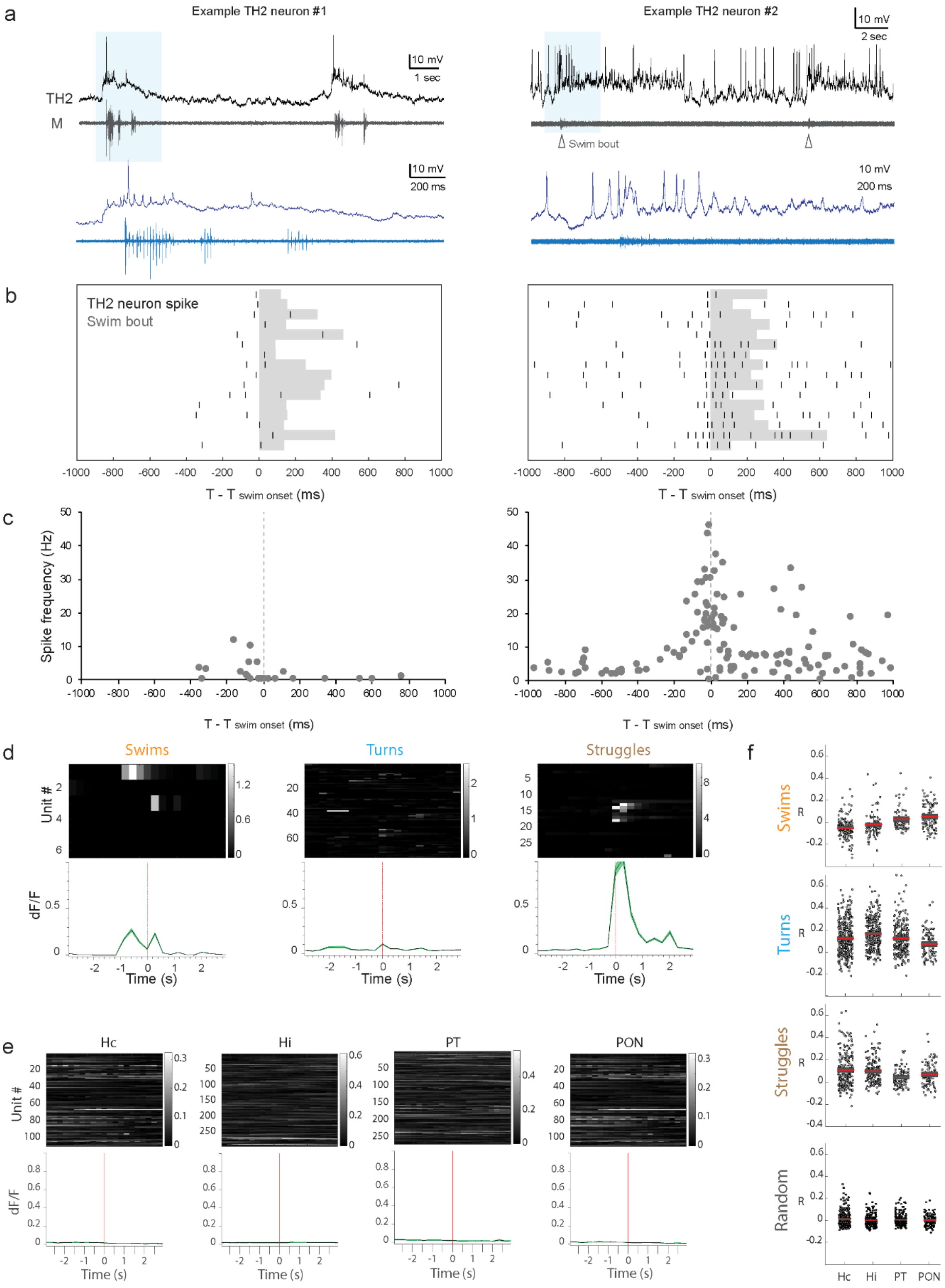

**Figure S2: Spontaneous, locomotion-associated activity of preoptic *th2*+ neurons**

(a) Example traces of whole-cell current clamp recordings from two *th2*+ neurons (left and right panels, "TH2" trace) with simultaneous EMG recordings ("M" trace) as a fictive metric of locomotion. The blue shaded regions are expanded in the lower panels.

(b) *th2*+ firing patterns during spontaneous swimming (n = 16 swim bouts for each *th2*+ neuron). Action potentials (black ticks) are aligned relative to swim bout initiation (grey shading).

(c) Instantaneous firing frequencies of the *th2*+ neurons shown in (b) aligned to the start of the associated swim bouts.

(d) Calcium traces of preoptic neurons acquired in *Tg(th2:gcamp5)* animals aligned to spontaneous behavior onset, separated by behavior type. Gray traces show the behavior-triggered average for individual units (mean dF/F). Green traces are mean +/- SEM across animals (N = 11 fish)

(e) Calcium traces of the *th2*+ neurons from figures 2 and 3 aligned to random frames of the calcium imaging movie. Gray traces show the behavior-triggered average for individual units (mean dF/F). Green traces are mean +/- SEM across animals (N = 20 fish). Activity in Hc, Hi, and PT neurons was recorded using Tg(*th2:gcamp5*), activity in PON was recorded using transient expressors of *th2:gcamp7s*.

(f) Pearson's R values from the regression of individual neurons and the behavior-specific regressors as well as a randomized control regressor, for which a number of randomized behavior onset times were chosen equal to the number of real behaviors occurring during the recording of the given neuron.

**Figure S3: Morphological registration to the z-brain**

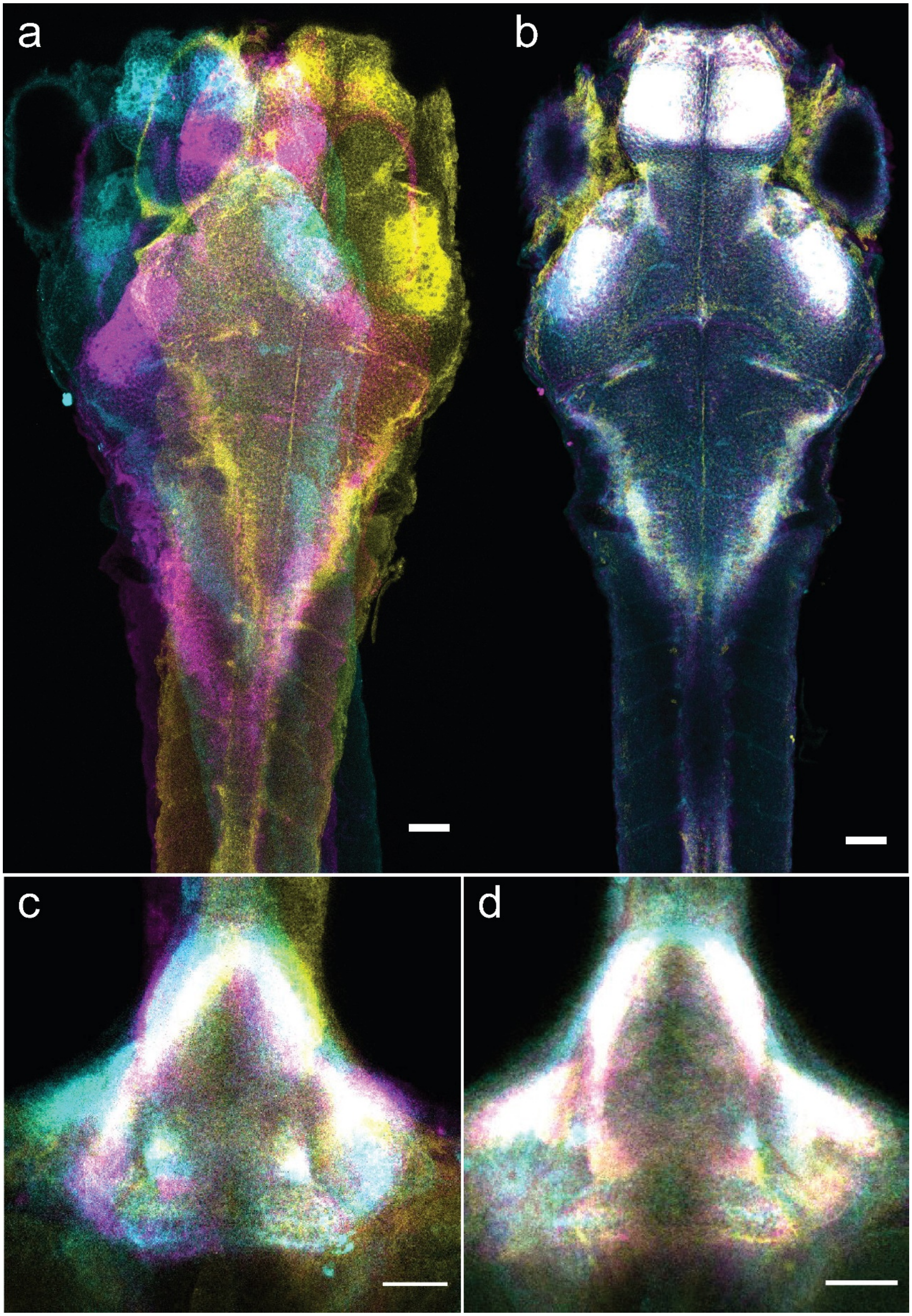

**Figure S3: Morphological registration to the z-brain**

(a) Confocal optical sections of total ERK (tERK) staining from three fish, pre-registration.

(b) Confocal optical sections from (a) after morphological registration to the z-brain reference atlas.

(c) Pre-registration confocal optical sections of the caudal hypothalamus (Hc).

(d) Post-registration confocal optical sections of Hc after morphological registration to the z-brain reference atlas.

### Figure S4: Estimation of sampling saturation for *th2*+ neuron projection types

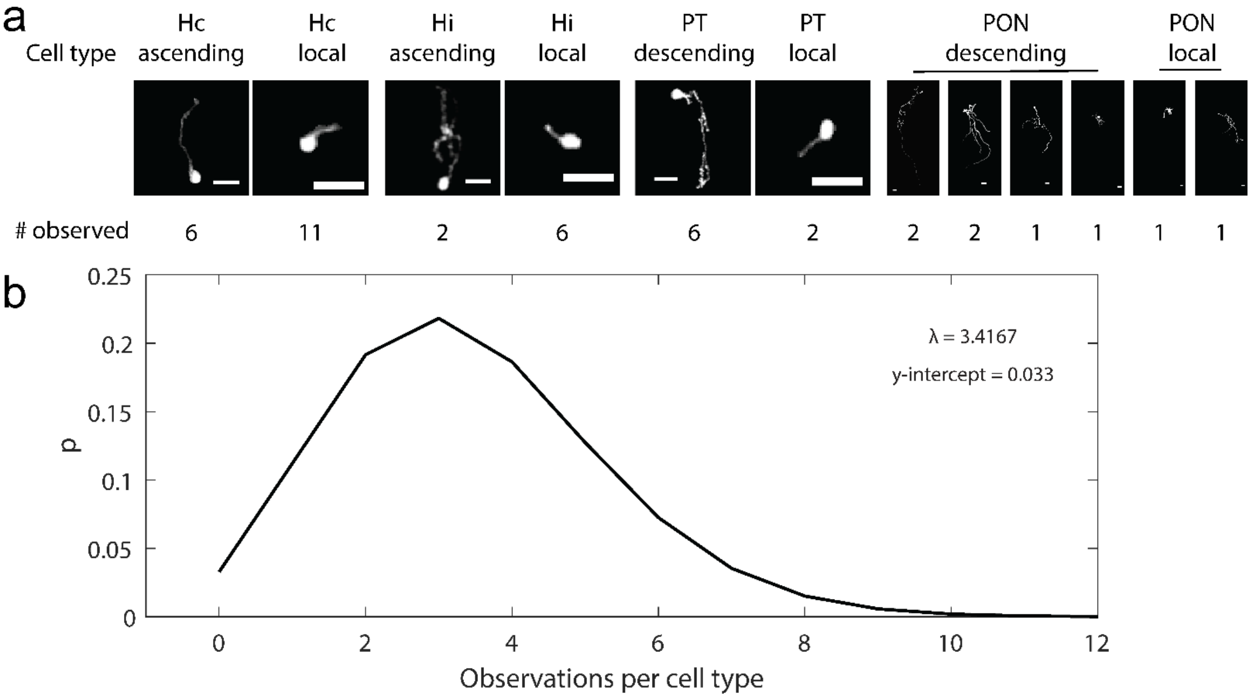

**Figure S4: Estimation of sampling saturation for *th2*+ neuron projection types**

(a) Maximum intensity projections showing examples of each *th2*+ cell type and the total number of cells observed.

(b) Poisson sampling distribution with lambda equal to the mean number of cells per cell type identified in our tracing experiments. Y-intercept represents the probability of 0 observations for a given cell type.

### Table S1: MAP-mapping reveals brain regions modulated by activity of *th2*+ neurons

| |
|---|
| Diencephalon - |
| Diencephalon - Caudal Hypothalamus |
| Diencephalon - Diffuse Nucleus of the Intermediate Hypothalamus |
| Diencephalon - Dorsal Thalamus |
| Diencephalon - Eminentia Thalami |
| Diencephalon - Habenula |
| Diencephalon - Hypothalamus - Caudal Hypothalamus Neural Cluster |
| Diencephalon - Hypothalamus Gad1b Cluster 3 Sparse |
| Diencephalon - Hypothalamus Vglut2 Cluster 6 |
| Diencephalon - Intermediate Hypothalamus |
| Diencephalon - Left Habenula Vglut2 Cluster |
| Diencephalon - Medial vglut2 cluster |
| Diencephalon - Olig2 Band |
| Diencephalon - Olig2 Band 2 |
| Diencephalon - Optic Chiasm |
| Diencephalon - Pituitary |
| Diencephalon - Posterior Tuberculum |
| Diencephalon - Postoptic Commissure |
| Diencephalon - Preoptic Area |
| Diencephalon - Right Habenula Vglut2 Cluster |
| Diencephalon - Rostral Hypothalamus |
| Diencephalon - Ventral Thalamus |
| Ganglia - Olfactory Epithelium |
| Ganglia - Statoacoustic Ganglion |
| Mesencephalon - |
| Mesencephalon - Medial Tectal Band |
| Mesencephalon - Oculomotor Nucleus nIII |
| Mesencephalon - Tectum Stratum Periventriculare |
| Mesencephalon - Tecum Neuropil |
| Mesencephalon - Tegmentum |
| Mesencephalon - Torus Semicircularis |
| Rhombencephalon - |
| Rhombencephalon - Cerebelluar-Vglut2 enriched areas |
| Rhombencephalon - Cerebellum |
| Rhombencephalon - Cerebellum Gad1b Enriched Areas |
| Rhombencephalon - Corpus Cerebelli |
| Rhombencephalon - Eminentia Granularis |
| Rhombencephalon - Gad1b Cluster 1 |
| Rhombencephalon - Gad1b Cluster 17 |
| Rhombencephalon - Gad1b Stripe 1 |
| Rhombencephalon - Glyt2 Cluster 1 |
| Rhombencephalon - Neuropil Region 5 |
| Rhombencephalon - Neuropil Region 6 |
| Rhombencephalon - Olig2 Stripe |
| Rhombencephalon - Olig2 enriched areas in cerebellum |
| Rhombencephalon - Ptf1a Stripe |
| Rhombencephalon - Raphe - Superior |
| Rhombencephalon - Rhombomere 1 |
| Rhombencephalon - Rhombomere 2 |
| Rhombencephalon - Rhombomere 3 |
| Rhombencephalon - Rhombomere 4 |
| Rhombencephalon - Rhombomere 5 |
| Rhombencephalon - Rhombomere 6 |
| Rhombencephalon - Rhombomere 7 |
| Rhombencephalon - S1181t Cluster |
| Rhombencephalon - Valvula Cerebelli |
| Rhombencephalon - Vglut2 Stripe 1 |
| Rhombencephalon - Vglut2 cluster 2 |
| Rhombencephalon - Vmat2 Stripe1 |
| Rhombencephalon - X Vagus motorneuron cluster |
| Spinal Cord |
| Spinal Cord - Gad1b Stripe 1 |
| Spinal Cord - Neuropil Region |
| Spinal Cord - Vglut2 Stripe 2 |
| Spinal Cord - Vglut2 Stripe 3 |
| Telencephalon - |
| Telencephalon - Olfactory Bulb |
| Telencephalon - Olig2 Cluster |
| Telencephalon - Optic Commissure |
| Telencephalon - Pallium |
| Telencephalon - Postoptic Commissure |
| Telencephalon - Subpallial Gad1b cluster |
| Telencephalon - Subpallium |
| Telencephalon - Vglut2 rind |

**Table S1: MAP-mapping reveals brain regions modulated by activity of *th2*+ neurons**

Names of annotated regions of the z-brain showing significantly increased pERK signal in Tg(*th2*:*gal4; uas:chr2-eyfp*) animals compared to non-expressing sibling controls after exposure to blue light. One-tailed Fischer's Exact test against the null hypothesis of 0 signal. N = 6 experiments, 15-20 animals per experiment, $p < 0.05$.

**Figure S5: *th2*+ projections target the nMLF**

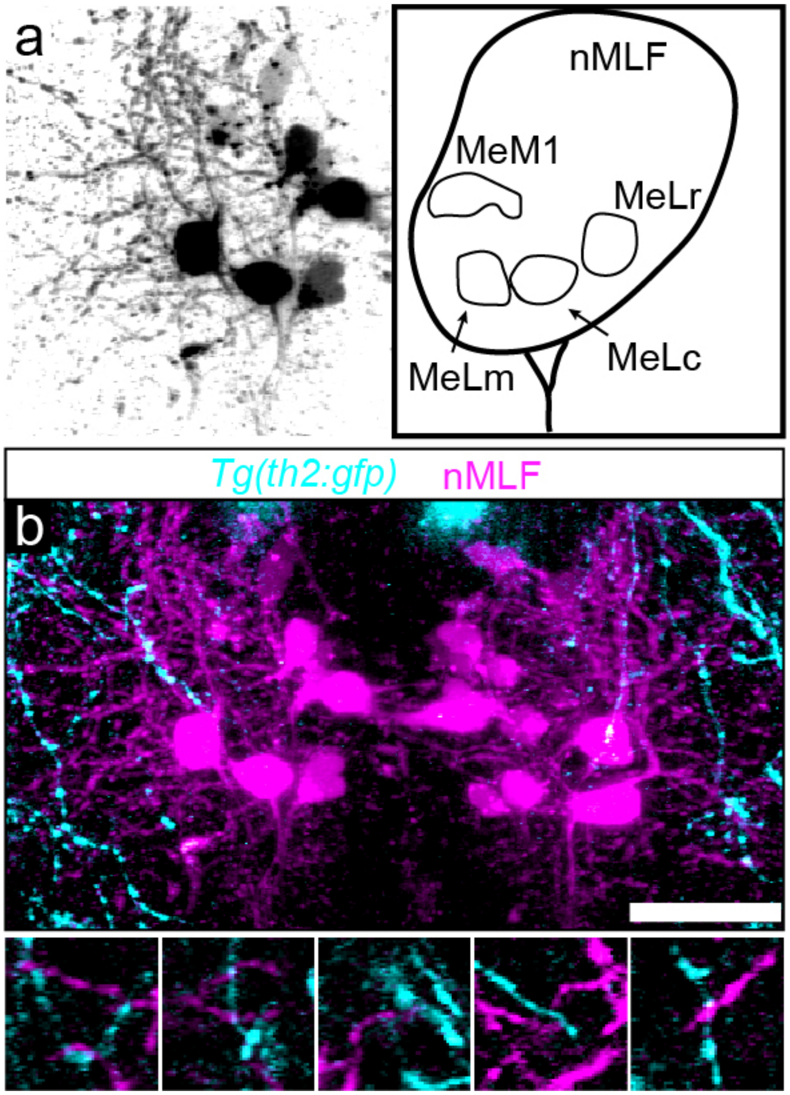

**Figure S5: *th2*+ projections target the nMLF**

(a) Maximum intensity projection of a two-photon z-stack showing neurons of the nMLF labelled by spinal backfill.

(b) Maximum intensity projection (top) and example single z-slices (bottom) showing *th2:gfp*+ terminals (cyan) in close apposition to nMLF soma and dendrites (magenta). Scale bar, 50 μm.

**Figure S7: Ablation of preoptic *th2*+ neurons reduces spontaneous swimming frequency**

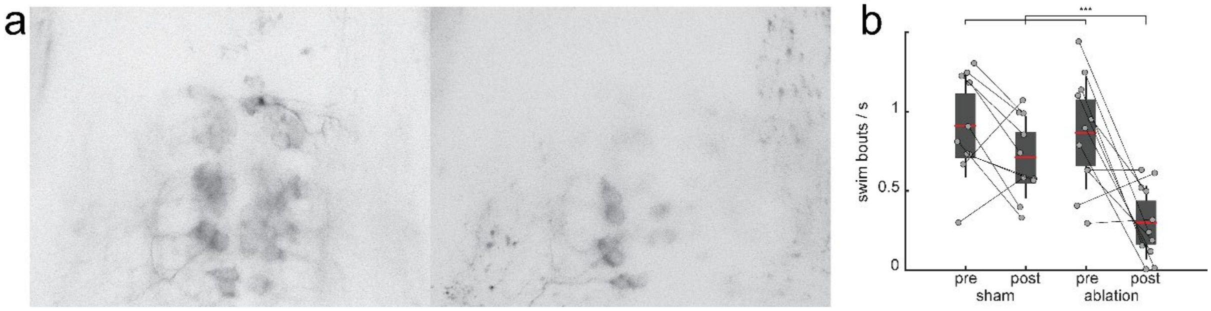

**Figure S7: Ablation of preoptic *th2*+ neurons reduces spontaneous swimming frequency**

(a) Maximum intensity projection of Tg(*th2*:*gfp-aequorin*)+ neurons in the PON, before and 24 hours after photoablation. 12.57 +/- 4.12 neurons were successfully ablated per fish (52.26 +/- 14.99%).

(b) Frequency of spontaneous swim bouts before and after sham ablation and before and after true ablation ($p < 0.0005$, Bonferonni-corrected ANOVA, $N = 10$ sham, 11 ablated).

**Movie S1: Behaviors identified by self-organizing map clustering**

Representative swim bouts of 6-8 dpf head-embedded larval zebrafish identified by self-organizing map clustering. Left, swim; middle, turn; right, struggle.  Acquisition, 500 fps; playback, 30 fps.

**Movie S2: Activity in Hc, Hi, and PT is highly correlated between clusters and with locomotor initiation**

Example calcium imaging experiment showing a head-embedded, tail-free Tg(*th2*:*gcamp5*) animal (top left) with tail angle measurement via eight equally spaced tracking points (bottom left). Calcium activity in Hc, Hi, and PT at right. Behavior acquisition, 500 fps; fluorescence acquisition, 3.44 fps; playback, 50 fps.
